# Metagenome-wide measurement of protein synthesis in the human fecal microbiota using MetaRibo-Seq

**DOI:** 10.1101/482430

**Authors:** Brayon J. Fremin, Ami S. Bhatt

## Abstract

The healthy human fecal microbiota is too diverse to comprehensively study with the current throughput of proteomic methods. Shotgun sequencing technologies allow for much more comprehensive profiling. Here, we develop and apply MetaRibo-Seq, a method for simultaneous ribosome profiling of multiple taxa within a complex bacterial community. This approach captures taxonomic diversity in fecal samples. As expected, the detected ribosome-bound transcripts are relatively enriched within coding regions and significantly correlate to detectable protein abundances. In a low diversity fecal sample, we show that MetaRibo-Seq is more strongly correlated than metatranscriptomic data to protein abundance. This significant correlation of metatranscriptomics and MetaRibo-Seq with protein levels is maintained, though with decreased strength as taxonomic diversity increases. Finally, we identify genes that are consistently regulated at the translational level across bacterial taxa within fecal communities. In conclusion, MetaRibo-Seq enables comprehensive translational profiling in complex bacterial communities for the first time.

## Introduction

There is great interest in determining the potential functions of the human fecal microbiota. To date, methods have excelled at describing the taxonomy of such communities; however, assigning and defining functions of the community of bacteria or individual organisms within these communities has been challenging^1^. An ideal method to study functions within a complex community would allow simultaneous enumeration of all of the proteins, lipids, and other macromolecules within the mixture. Unfortunately, this is not feasible with current technologies. While 10^2^ to 10^4^ proteins can be simultaneously quantified with metaproteomics^2^, it is challenging to obtain accurate measurements of the full array of bacterial proteins that likely exist in human fecal samples, estimated to be 10^7^ to 10^8^ proteins^3^. Thus, current proteomic methods lack the dynamic range required to comprehensively study the human fecal microbiota^4^.

Given the challenges in direct protein measurement and need for databases of protein sequences, some have focused on enumerating the gene content of a community to determine the potential function; indeed, progress has been made in predicting genes present within a metagenome^5^. However, the presence of a gene in a complex bacterial community does not imply that the gene is transcribed or translated. Acknowledging this limitation, recent work has demonstrated that differential transcription of bacterial genes can be used to derive biologically meaningful insights^6,7,8,9^. Yet, we still have a very limited understanding of regulation occurring post-transcriptionally in the human fecal microbiota.

In contrast to transcriptomic profiling, ribosome profiling (Ribo-Seq) is a method that quantifies protein synthesis^10^. In eukaryotes, Ribo-Seq generally correlates more strongly to protein abundance than transcriptomics^11,12,13^; this correlation has not been described in bacteria. Bacterial ribosome profiling studies have been performed in model organisms such as *Escherichia coli* and *Bacillus subtilis,* with minor modifications from the eukaryotic protocols, such as using chloramphenicol to inhibit translation and micrococcal nuclease (MNase) to enrich for ribosome footprints^13,14,15,16,17^. These methodological modifications enable a high-throughput snapshot of translation, but often compromise the ability to achieve codon-level resolution^11^.

In bacteria, many genes are regulated at the translational level. For example, genes involved in translation itself are known to be regulated at a translational level via feedback mechanisms^18,19,20,21,22^. Translational regulation is critical for generating proteins at the correct stoichiometry for many protein complexes. For example, the multiprotein complex that forms bacterial ATP synthase has multiple genes whose translation is regulated; the stoichiometry of this complex is best predicted by Ribo-Seq^13,17^. This even extends to pathway-specific enzyme stoichiometry in which protein synthesis remains conserved as compensation for transcript abundance and architecture divergence across taxa17. Moreover, specific translational regulation has been extensively observed upon a variety of perturbations to bacteria^23,24,25,26,27,28,29^. For example, bacteria employ translational quality control and regulation of amino acid biosynthesis in response to amino acid stress^30^. Thus, translation is a conserved, critical, dynamic, and regulated process in bacteria. However, this level of regulation has thus far been overlooked in mixed bacterial communities. Previous studies of protein synthesis in bacteria have been restricted to pure large-scale cultures (media and RNA inputs up to liters and milligrams, respectively). To date, studying translational regulation in mixed communities or in culture-free contexts has been hindered by low extraction yield, low purity, and the lack of informatic frameworks to study organisms without reference genomes. Consequently, we have a very limited understanding of how widespread bacterial translational regulation may be outside of cultured model organisms.

In this work, we develop a method that allows for simultaneous ribosome profiling in a complex community of bacteria without the need for a large-scale, purified cultures. With three lines of evidence, we confirm that MetaRibo-Seq effectively enables translation to be studied in the fecal microbiota. First, the signal consists of footprints that capture the taxonomic diversity of metagenomics, while being locally enriched within coding regions. We identify most enrichment at gene start and stop codons, characteristic of chloramphenicol-treated ribosome profiling^31^. Second, ribosome footprint densities significantly correlate to detectable protein abundances and are significantly enriched in signal for these abundant proteins in these complex bacterial communities. In low diversity human fecal samples, we show that MetaRibo-Seq better correlates with protein abundance and *E. coli* ATP synthase stoichiometry than transcriptomics. Third, biological processes known to be translationally regulated, such as translation itself, are consistently detected as such across multiple samples and taxa. We catalog tens of thousands of genes with evidence of translational regulation in fecal samples across diverse taxa, providing a widespread view of consistent, bacterial translational regulation in these systems. Overall, we show that MetaRibo-Seq facilitates metagenome-wide measurement of bacterial protein synthesis across taxa directly in fecal samples.

## Results

### The MetaRibo-Seq workflow

MetaRibo-Seq enables sequencing of ribosome-protected footprints directly from human fecal samples (see Methods, Figure 1A). First, we show ribosome profiling can be performed on frozen fecal samples stored in RNAlater^32, 6^ (Ambion), an RNA preserving solution. Unlike some existing protocols^33,16^, our ribosome profiling protocol first introduces chloramphenicol during lysis. Bead beating lysis is performed to also lyse diverse Gram-positive bacteria^34^. An ethanol precipitation step post-lysis is introduced to both filter out fecal debris and concentrate RNA and any complexes bound. This has been demonstrated to effectively precipitate ribosomes^35^. MNase treatment is performed on an extremely crude purification of nucleic acids and complexes. Ribosome profiling performed here uses nearly an order of magnitude less RNA and MNase than isolate protocols typically use (see Methods)^16, 33^. After high-quality footprints are reliably generated using these methods, ribosome profiling converges to isolate protocols to purify monosomes and prepare libraries^33^. MetaRibo-Seq overcome challenges of sample storage, input requirement, bacterial purity, and uniform lysis to generate high quality RNA footprints from fecal samples.

**Figure 1.**
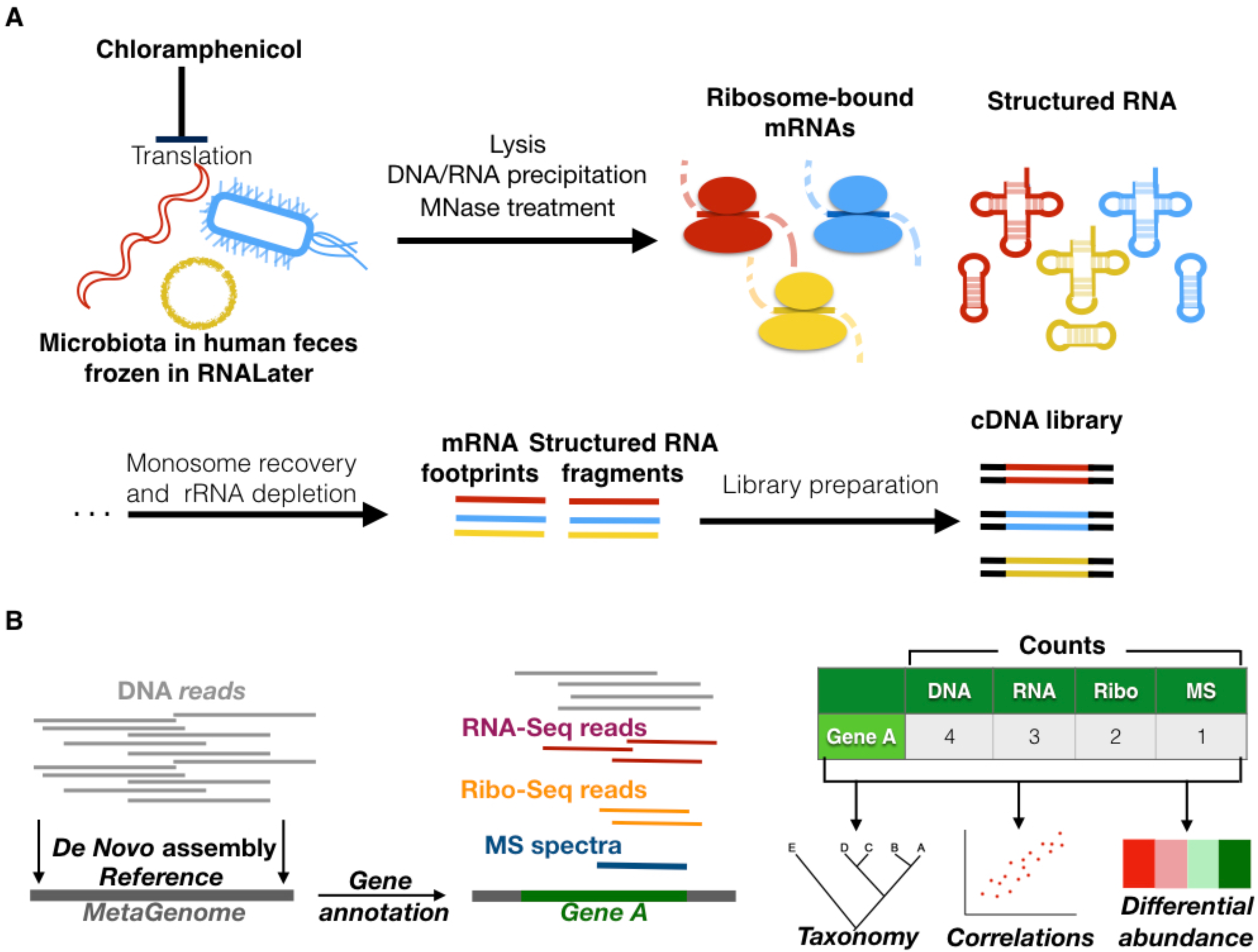
Workflow of ribosome profiling. (A) Experimental workflow of MetaRibo-Seq. Chloramphenicol halts translation, the bacterial community is lysed, MNase is used to create footprints, and footprints are converted to sequencing libraries. (B) Computational workflow of the multi-omics approach. *De novo* assemblies are created and annotated, predicted genes are quantified at multi-omic levels, and taxonomy, correlations, and differential abundance are determined from these results.

Computationally, dealing with short reads and poor or incomplete reference genomes is challenging. To overcome these challenges, we use a *de novo* approach to build references, annotate genes, and map reads to those references (see Methods, Figure 1B). Mapping metrics to *de novo* references are provided (Table S1). We require perfect, unique matches of these ribosome footprints to references to ensure proper mapping. Multi-mapping varies sample to sample, ranging from 6.95 to 26.69 percent. We find that 4.1 - 10.4 percent of mapped reads from RNA technologies performed correspond to predicted coding regions. Given variable amounts of diversity and heterogeneity in any given sample, mapping statistics will vary sample to sample.

### MetaRibo-Seq signal retains taxonomic diversity in human fecal samples

MetaRibo-Seq captures taxonomic bacterial diversity in human fecal samples via ribosome-protected footprints. We perform MetaRibo-Seq on four diverse fecal samples. Sample A is from a healthy individual. Sample B is from a patient with a hematological disorder who is undergoing treatment. Sample C is from a patient with a solid malignancy who is undergoing treatment. Sample D is from a patient with Alzheimer’s disease. We also perform MetaRibo-Seq on a low diversity fecal sample from a patient with a hematological disorder who is undergoing antibiotic treatment with metronidazole – Sample E. Metagenomic reads are subjected to *de novo* assembly and gene prediction and annotation for each sample (see Methods). These assemblies and gene predictions are provided (NCBI BioProject ####). Taxonomic differences at the genus level exist between technologies across samples, though most abundant taxa are largely consistent across technologies (Figure 2A). Shannon diversity is also concordant between technologies, including MetaRibo-Seq (Figure 2B). Thus we conclude that MetaRibo-Seq signal faithfully recapitulates the diversity of organisms present in the mixed bacterial communities.

**Figure 2.**
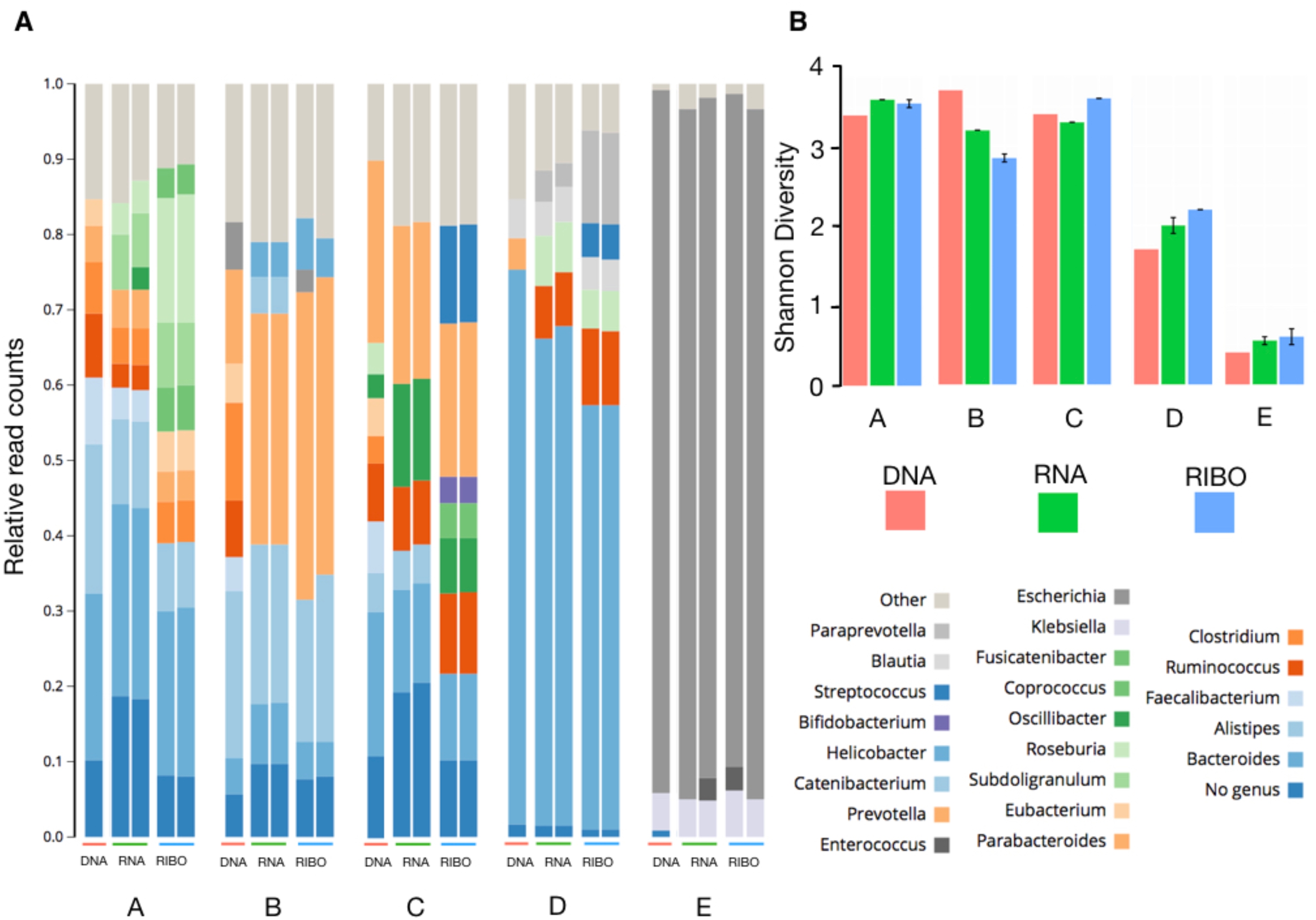
MetaRibo-Seq signal captures diversity in a metagenomic context. (A) Genus-level classifications of all sequencing technologies performed on Samples A, B, C, D, and E. Replicates for metatranscriptomics and MetaRibo-Seq are shown for reproducibility. Taxa represented below three percent are grouped into “Other” for visual purposes. (B) Shannon diversities across technologies for these samples are displayed.

### MetaRibo-Seq signal is characteristic of bacterial ribosome profiling

We find that MetaRibo-Seq signal is locally enriched within coding regions. We show average signal across all coding predictions and flanking regions for Samples A, B, C, and D (Figure 3A-D). We visualize strong signal corresponding to predicted ORFs with pronounced signal drop off outside of the start and stop codons for samples A through D (Figure 3A-D). Start and stop codons represent the strongest signal. Surprisingly, MetaRibo-Seq also shows some weak signs of overall codon resolution (Figure S1). In a more targeted analysis, MetaRibo-Seq can achieve stronger codon resolution of ribosomes in common genera. For Sample A and Sample B, assembled contigs of several shared genera are classified and binned appropriately (see Methods). Triplet periodicity is observed across footprint lengths in Bacteroides (Figure S2A), Faecalibacterium (Figure S2B), and Alistipes (Figure S2C). Based purely on raw signal, these findings collectively suggest that MetaRibo-Seq is capturing ribosome-bound footprints as expected.

**Figure 3.**
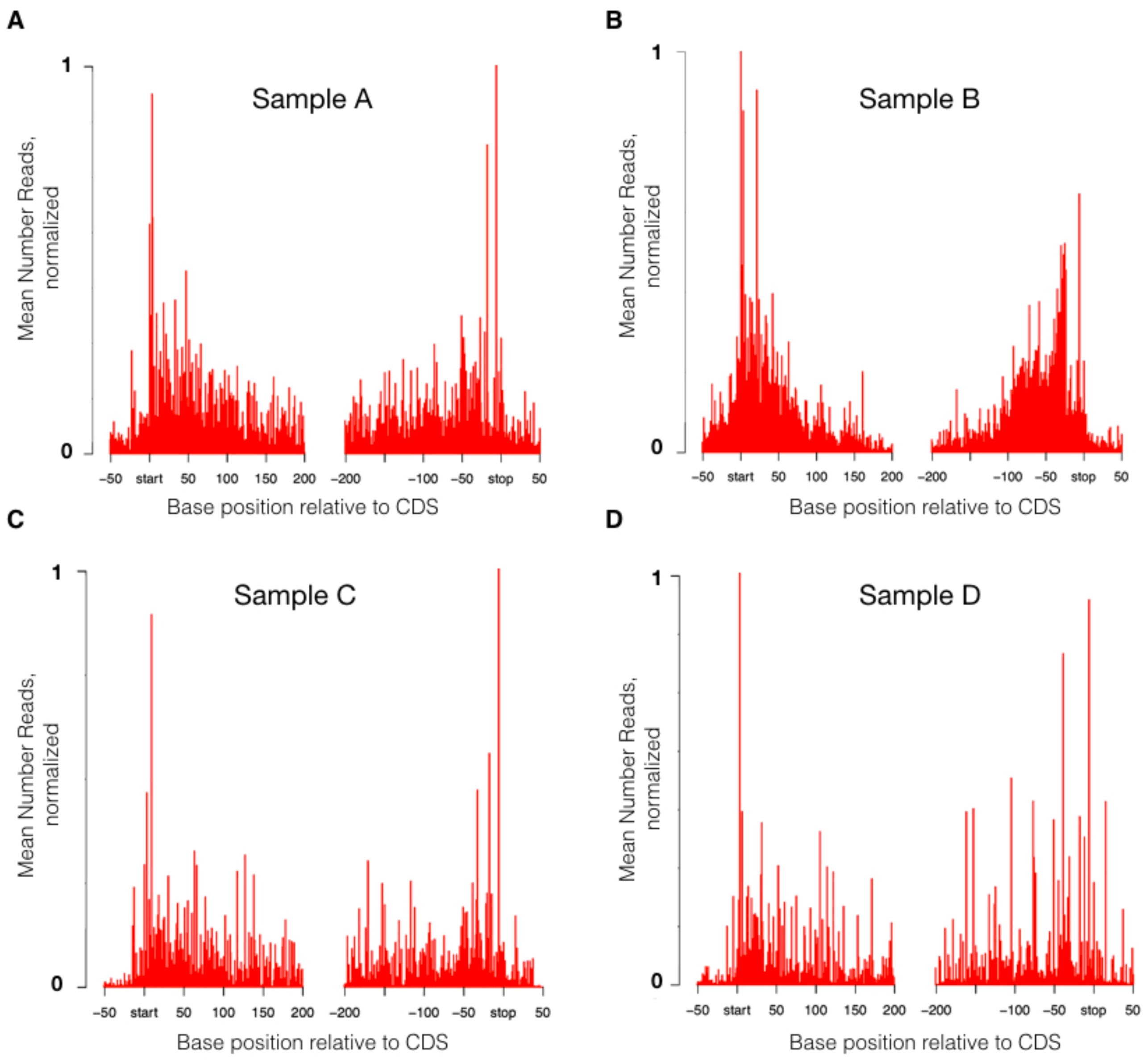
MetaRibo-Seq signal is characteristic of chloramphenicol-treated ribosome profiling in bacteria across diverse samples. (A-D) Average MetaRibo-Seq signal across genes and flanking regions for Sample A, B, C, and D, respectively. Every predicted open reading frame containing at least 10 reads are included in the analysis.

### MetaRibo-Seq outperforms metatranscriptomics as a proxy for protein abundance in a low diversity fecal sample

In Sample E, we identify strong correlations of metatranscriptomics and MetaRibo-Seq to metaproteomics. First, Sample E is dominated by *E coli*. Interestingly, this *E. coli* is later isolated from the blood of the patient with no single nucleotide variants compared to fecal *E. coli*^36^. This blood isolate has been sequenced and represents our isolate reference for downstream analyses. In a separate study demonstrating strain identity, this patient is denoted as Patient 3, and Sample E in this study is specifically 27 days prior to bacteremia^36^. For the 1503 genes that were proteomically-detected in Sample E, we show genus-level representation for metatranscriptomics, MetaRibo-Seq, and metaproteomics (Figure 4A). Of note, 812 of these proteins belong to *E. coli*, while the remaining 691 belong to other taxa. There is greater representation of proteins predicated to arise from Klebsiella and Enterococcus than transcripts or MetaRibo-Seq signal (Figure 4A). We show a Pearson correlation of 0.46 between MetaRibo-Seq and metaproteomics across the 1503 detected proteins from the metagenomic Sample E (Figure 4B). The Pearson correlation is 0.64 when only considering the 928 detected proteins from the *E. coli* isolate reference (Figure 4C). We display correlations between metatranscriptomics, MetaRibo-Seq, and metaproteomics (Figure 5D). Same technology correlations between the 812 identical protein predictions from the metagenomic and isolate *E. coli* analyses all retain Pearson correlations of 0.99. Correlations are weaker in the metagenomic context specifically due to relatively poorer predictions upon addition of the 691 proteins that do not belong to *E. coli*. No previous study to our knowledge provides a correlation between ribosome profiling and proteomics in *E. coli* or any other bacteria; however, this correlation of 0.64 is stronger than any cited correlation between transcriptomics and proteomics in isolated, cultured, model *E. coli*^37^. ATP synthase is a well-characterized complex in terms of stoichiometry in *E. coli* (Figure 4E). We show that MetaRibo-Seq signal better correlates with known ATP synthase stoichiometry than transcriptomics, as expected (Figure 4F-G). Among sequencing technologies, MetaRibo-Seq serves as a better proxy for protein levels and ATP synthase stoichiometry in Sample E *E. coli*.

**Figure 4.**
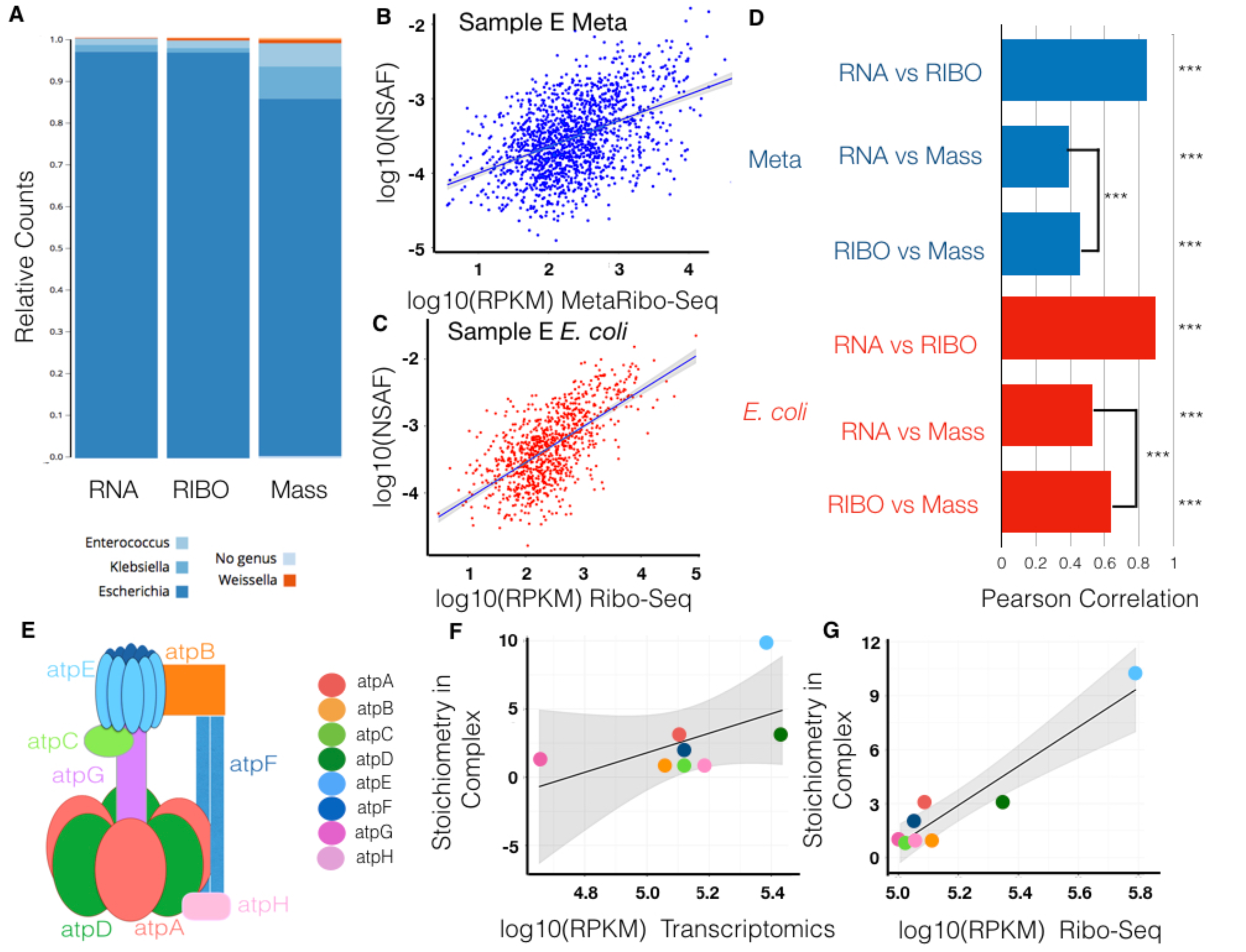
In a low diversity fecal sample, MetaRibo-Seq is a significantly better predictor than metatranscriptomics of protein abundance and ATP synthase stoichiometry in *E. coli*. (A) In Sample E, there are 1503 proteomically-detectable genes. Only focusing on these genes, we taxonomically classify the entire gene in equal number to the reads (for metatranscriptomics and MetaRibo-Seq) or spectral counts (for metaproteomics) assigned to it (see Methods). (B) Scatterplot of MetaRibo-Seq RPKM and metaproteomics NSAF log-scaled for these 1503 genes. (C) Scatterplot of MetaRibo-Seq RPKM and metaproteomics NSAF log-scaled only for the 928 proteomically-detected genes predicted from isolate *E. coli*. (D) Pearson correlations for pairwise comparisons across technologies. Blue bars indicate Pearson correlations pertaining to the entire metagenomic Sample E. The correlations between metatranscriptomics vs. MetaRibo-Seq, metatranscriptomics vs. metaproteomics, and MetaRibo-Seq vs. metaproteomics are 0.85, 0.39, and 0.46, respectively. All are significant (p value < 2^-16^). MetaRibo-Seq is a significantly better predictor of protein levels with Zou’s^58^ 95 % confidence interval between -0.0917 to -0.0487. Red bars indicate Pearson correlations pertaining to the isolated *E. coli* in the sample. The correlations between metatranscriptomics vs. MetaRibo-Seq, metatranscriptomics vs. metaproteomics, and MetaRibo-Seq vs. metaproteomics are 0.90, 0.53, and 0.64, respectively. All are significant (p value < 2^-16^). MetaRibo-Seq is a significantly better predictor of protein levels with Zou’s^58^ 95 % confidence interval between -0.1352 to -0.0868. (E) F0F1 ATP synthase in *E. coli* forms with a specific stoichiometry as visualized. (F) Correlation between log-scaled metatranscriptomics RPKM and expected ATP synthase stoichiometry in complex. The Pearson correlation is 0.56 (p value = 0.1475) (G) Correlation between log-scaled MetaRibo-Seq RPKM and expected ATP synthase stoichiometry in complex. The Pearson correlation is 0.92 (p value = 0.0012).

**Figure 5.**
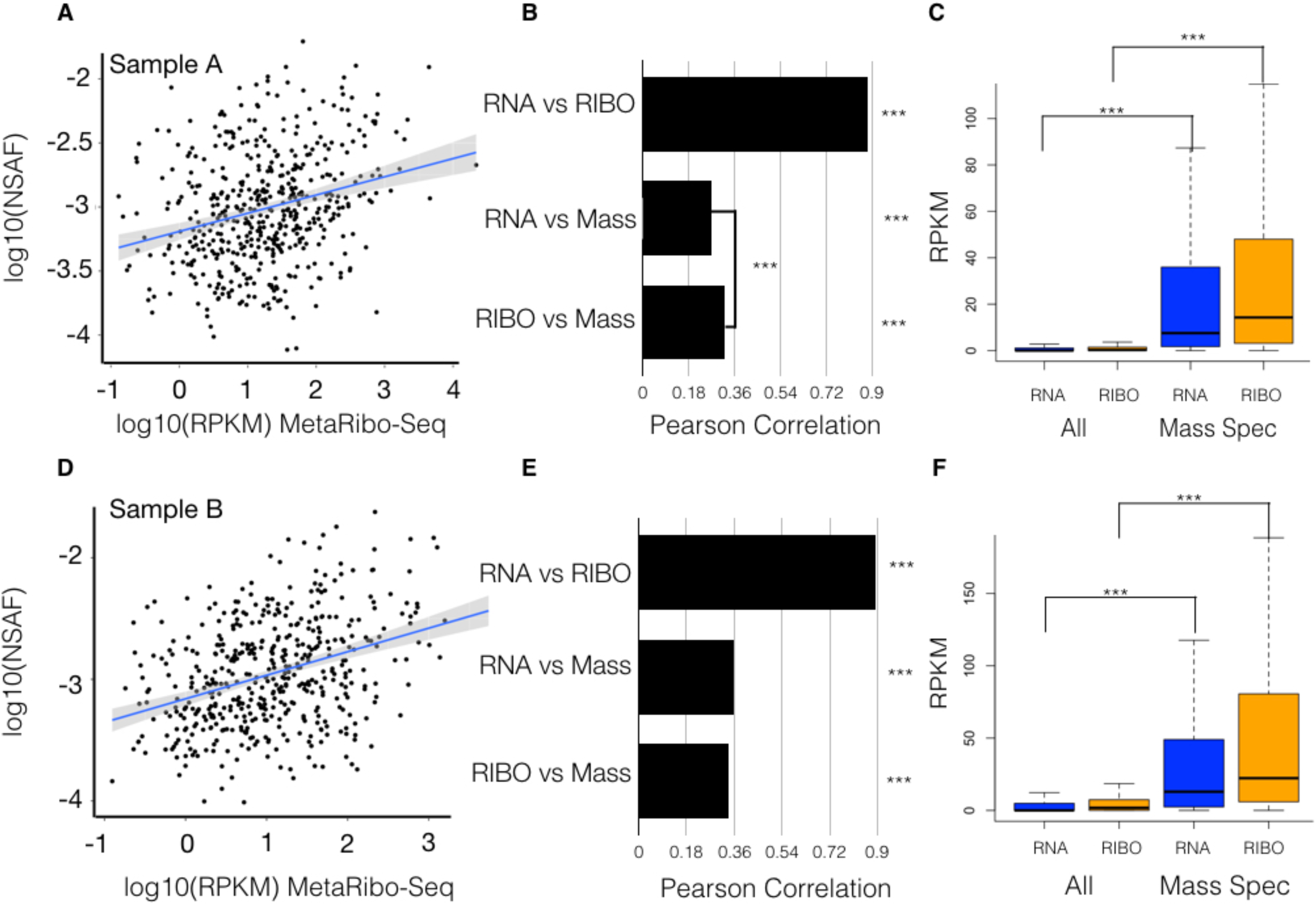
MetaRibo-Seq signal significantly correlates to metaproteomics and is enriched for these products. (A) For Sample A, scatterplot of MetaRibo-Seq RPKM (Reads per Kilobase Million) and metaproteomics NSAF (Normalized Spectral Abundance Factor) both log10-scaled. 497 genes are displayed. (B) Pairwise Pearson correlations between log-scaled metatranscriptomics RPKM, MetaRibo-Seq RPKM, and metaproteomics NSAF for these 497 genes. Pearson correlations are 0.88, 0.26, and 0.32 for metatranscriptomics vs. MetaRibo-Seq, metatranscriptomics vs. metaproteomics, and MetaRibo-Seq vs. metaproteomics, respectively. All are significant (p value < 2^-16^). MetaRibo-Seq is a significantly better predictor of protein levels than metatranscriptomics for these proteins in Sample A with a Zou’s^58^ 95 % confidence interval between -0.1322 and -0.0480. (C) Metatranscriptomics and MetaRibo-Seq RPKM for all predicted genes compared to those detected by metaproteomics. Both metatranscriptomic and MetaRibo-Seq signal for proteomically-detected proteins are significantly enriched (p value < 2^-16^). (D) For Sample B, scatterplot of MetaRibo-Seq RPKM and metaproteomics NSAF both log10-scaled. 497 genes are displayed. (E) Pairwise Pearson correlations between log-scaled metatranscriptomics RPKM, MetaRibo-Seq RPKM, and metaproteomics NSAF for these 480 genes. Pearson correlations are 0.89, 0.36, and 0.34 for metatranscriptomics vs. MetaRibo-Seq, metatranscriptomics vs. metaproteomics, and MetaRibo-Seq vs. metaproteomics, respectively. All are significant (p value < 2^-16^). (F) Metatranscriptomics and MetaRibo-Seq RPKM for all predicted genes compared to those detected by metaproteomics. Both metatranscriptomic and MetaRibo-Seq signal for proteomically-detected proteins are significantly enriched (p value < 2^-16^).

### MetaRibo-Seq signal significantly correlates to protein abundance and is enriched in these proteins in mixed bacterial communities

We find that MetaRibo-Seq significantly correlates to protein abundances as measured by shotgun metaproteomics. In Sample A, we detect 497 proteins. We measure a significant Pearson correlation of 0.32 between MetaRibo-Seq and metaproteomics (Figure 5A). MetaRibo-Seq is better correlated with protein abundance than metatranscriptomics in Sample A (Figure 5B). As expected, both metatranscriptomics and MetaRibo-Seq are significantly enriched in signal for these 497 proteomically-detected genes (Figure 5C). In Sample B, we detect 480 proteins and measure a Pearson correlation of 0.34 between MetaRibo-Seq and metaproteomics (Figure 5D). There is no significant difference in protein abundance prediction between metatranscriptomics and MetaRibo-Seq in Sample B (Figure 5E). Metaranscriptomics and MetaRibo-Seq are similarly enriched in signal for these proteins in Sample B (Figure 5F). These proteins detected by mass spectrometry represent highly abundant bacterial proteins in the fecal samples. These findings suggest that MetaRibo-Seq correlates well with highly abundant protein levels in mixed bacterial communities, and that MetaRibo-Seq signal may thus serve as a surrogate for protein abundance in the study of complex bacterial communities.

MetaRibo-Seq signal characteristics and predictive power of protein abundance suggests that it may also prove useful in predicting proteins in taxonomically diverse fecal samples. As a preliminary demonstration of this, we predict small proteins using Prodigal^38^ with decreased length cutoff (see Methods). We show a histogram of the number of small predictions (20-29 amino acids) and the number of those predictions with MetaRibo-Seq RPKM greater than 0.5 across samples (Figure S4A). Due to metaproteomic limitations, we are unable to validate these proteins directly. However, we can use a comparative genomic approach to identify clusters of small proteins, all with evidence of translation, that also possess evolutionary signatures indicative of coding regions. We cluster proteins at 70 percent amino acid identity (see Methods). We discover 21 clusters (with at least 4 members) of small proteins across Samples A, B, C, and D that contain both translational evidence among all members in the cluster and significant coding signatures determined via RNAcode^39^ (Figure S3B). We also show greater protein conservation among predictions with MetaRibo-Seq signal than by random chance (Figure S3D). Translational evidence of small proteins in diverse fecal samples decreases the number of predictions to a more conserved subset, suggesting it may be useful in gene prediction.

### Consistent translational regulation is observed across samples and taxa

By contrasting metatranscriptomics with MetaRibo-Seq, we identify translationally regulated genes in fecal samples. This provides a widespread view of genes that are consistently translationally regulated within these systems. For Samples A, B, C, and D individually, we show the number of gene predictions and significantly translationally regulated genes (Figure 6A). We detect reasonable DESeq2 model fits in comparing between technologies, as shown by the dispersion plots for these analyses for each sample (Figure S4). These significantly translationally regulated genes are clustered at 70 percent amino acid identity for each sample (see Methods, Figure 6B). In a combined analysis (Figure 6C), we define any cluster containing five or more sequences as consistently translationally regulated. The representative sequences for all of these consistently translationally regulated clusters are assigned GO terms with Blast2GO^40^ (see Methods). The top 10 most common biological process associated GO terms are displayed, with translation being the top hit (Figure 6D). These sequences and clusters are provided for reference (File S1). Across samples, we catalog 42,267 differentially translated genes and 607 consistently translationally regulated gene clusters in these fecal samples, many of which are involved in expected processes, like translation.

**Figure 6.**
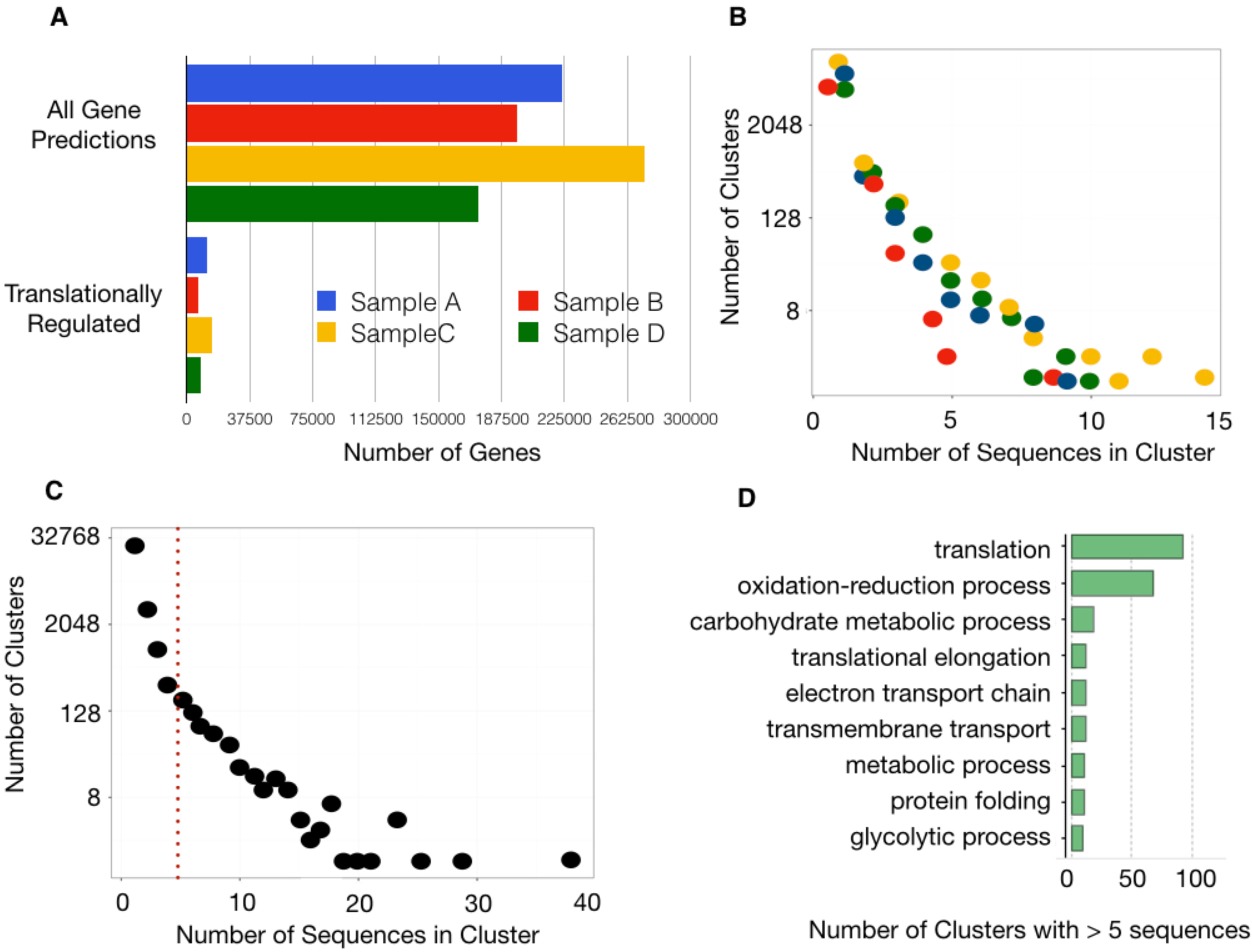
Genes that are consistently translationally regulated emerged across samples and taxa. (A) For Samples, A, B, C, and D, we show the number of total gene predictions via Prokka^41^. We performed DESeq2^60^ on these individually, comparing metatranscriptomics to MetaRibo-Seq. We show the number of those genes identified as translationally (absolute value(log2 fold change) > 1 and FDR < 0.05). We predict 223630, 196683, 272895, and 173624 genes from Sample A, B, C, and D, respectively. Among these, 11872, 6580, 15188, and 8647, respectively, are called significant translationally regulated genes. (B) Significant translationally regulated genes are converted to proteins and clustered at 70 percent amino acid identity (see Methods). The number of clusters with specific numbers of sequences are displayed, jittered and color-coded for each sample. (C) The number of clusters with specific numbers of sequences combined across Samples A, B, C, and D (D) If the combined cluster contained at least 5 sequences, this gene is considered consistently translationally regulated. 607 clusters met this requirement. For these clusters, the representative sequence (see Methods) is selected to represent the entire cluster. These representatives were input into Blast2GO to assign GO terms based on protein sequence.

Other approaches, albeit more biased to known gene annotations, are to rely on Prokka^41^ annotations. We count the number of times a gene symbol appears as differentially translated (Figure S5) and input those that appear at least five times into a GO Analysis (see Methods, Figure S6). Translation remains the top hit. For Samples A, B, C, and D, we provide GO analysis results of differential genes for each sample individually (Table S2). As a more pathway-oriented analysis, we also determine overrepresentation of pathways among globally differentially translated enzymes based on EC numbers assigned by Prokka^41^ (see Methods, Table S3). Amino acid biosynthesis is among the most consistent top hits across samples (Table S3). Thus several approaches lead to expected conclusions of pathways and processes that are translationally regulated.

## Discussion

One of the major limitations in advancing the functional knowledge of microbial communities is an inability to measure the macromolecular output of a given community in an unbiased manner. For example, until now, we have been unable to study fecal bacterial communities, or any *in vivo* system of bacteria, at the level of protein synthesis. Previous approaches have required *in vitro* growth of large, purified cultures; this limits both throughput and the diversity of the sample that can be studied. Here, we introduce a new method, MetaRibo-Seq, and provide evidence that this method enables the human fecal microbiota to be studied at a translational level. In five taxonomically varied samples from human subjects with variable health status, we show MetaRibo-Seq signal retains the taxonomic diversity of the samples. The signal itself is as expected for a chloramphenicol-treated ribosome profiling library – including local enrichment within coding regions and greater enrichment across the start and stop codons of genes. This suggests we have a way to measure bacterial protein synthesis *in vivo* for the first time.

To conduct a fair comparison, we perform MetaRibo-Seq on diverse samples but also a lower diversity fecal sample (Sample E) so that more representative protein quantifications of select taxa are achievable. In taxonomically diverse stool samples, MetaRibo-Seq is comparably predictive to protein abundance as metatranscriptomics. However, we show it can be significantly more predictive in a lower diversity scenario. We show that the addition of lower abundant taxa weaken overall correlations in mixed communities. Biologically, metatranscriptomics and MetaRibo-Seq are snapshots of gene transcripts or proteins synthesized, respectively, not direct measurement of the proteins that currently exists. Moreover, significant post-translational differences between taxa likely exist. Technically, it becomes challenging to obtain accurate protein abundances for lowly abundant taxa and proteins in a sample, making such correlations to protein abundance themselves less representative. We conclude that while MetaRibo-Seq can outperform metatranscriptomics within a highly abundant organism, this effect is diminished, perhaps both for biological and technical reasons, when considering all taxa together in diverse communities.

There are several limitations to MetaRibo-Seq. First, MetaRibo-Seq does not include steps to degrade RNAs with secondary structure. This is a common issue for ribosome profiling protocols but exacerbated in this *de novo*, low input context^10, 13^. Though targeted approaches for specific bacteria have been successful for tRNA depletion,^26^ an untargeted approach, which would be necessary here, has yet to be implemented in literature. Utilization of sucrose density gradients instead of size-exclusion chromatography columns may also prove valuable in removing various structured RNAs; however, downstream ribosome profiling input requirements likely make this challenging to address. This limitation may, however, enable other types of investigation; for example, signal corresponding to structured RNAs will likely be useful to predict novel structured RNAs in non-coding regions. Experimental modifications will also likely improve these limitations.

We anticipate that MetaRibo-Seq will enable a clearer functional view of the fecal microbiota. With increasing use of metatranscriptomics, we envision that MetaRibo-Seq will be applied to various disease states to better probe the microbiota and its functions. We especially anticipate MetaRibo-Seq will be used longitudinally to study translational regulation of genomically-stable, clinically-relevant taxa. MetaRibo-Seq signal provides unique features like enrichment within coding regions, greater enrichment at the start and end of genes, and, as we show, some signs of codon-level resolution in some taxa. Future work will likely include using these features for gene prediction, which is particularly challenging when studying metagenomic samples and small proteins^42^. To validate such predictions, significant methods development and improvements in metaproteomics will be needed. With direct proteomic evidence often unattainable, coding potential, translational evidence, and conservation among predictions present themselves as the strongest lines of evidence proteins, especially small proteins, exists in the fecal microbiota (Figure S6 and File S2). We also expect MetaRibo-Seq to be applied to other culture-free conditions, perhaps requiring other modifications. Overall, we show that translation can be comprehensively studied in mixed bacterial communities in a culture-free manner. This method also sheds light on consistently translationally regulated genes *in vivo* in a comprehensive, metagenome-wide analysis.

## Materials and Methods

### Subject Recruitment

MetaRibo-Seq was performed on fecal samples from individuals from a variety of health states. Informed consent was obtained for all participants. None of the participants received bacterial translation inhibitors. All subjects were recruited at Stanford University as a part of one of three IRB-approved protocols for tissue biobanking and clinical metadata collection (PIs: Dr. Ami Bhatt, Dr. Victor Henderson, Dr. David Miklos).

### Fecal Samples Storage

Stool was immediately stored in 2 mL cryovials and frozen at -80 °C. Stool was not thawed until lysis. For RNA extraction applications, 1.3 grams of fecal samples were preserved in 700 μL of RNALater (Ambion) at -80 °C.

### Cell Lysis for Metatranscriptomics and MetaRibo-Seq

Stool (150 mg) was suspended in 600 μL Qiagen RLT lysis buffer supplemented with one percent beta-mercaptoethanol and 0.3 U/μL Superase-In (Invitrogen). For MetaRibo-Seq lysis, 1.55 mM of chloramphenicol was also added to this lysis solution, and the solution was incubated at room temperature for 5 minutes. The suspension was subjected to bead beating for 3 minutes using 1.0 mm Zirconia/Silica beads. This was performed with a MiniBeadBeater-16, Model 607. The lysed solution was centrifuged at room temperature for 3 minutes at 21,000 x g to pellet cellular debris, and the supernatant was extracted to 2 mL tubes.

### Metagenomics

DNA was extracted from fecal samples with DNA Stool Mini Kit (Qiagen) using manufacture protocols. Samples were exposed to bead beating for 3 minutes. 1 ng of DNA was used to create Nextera XT libraries according to manufacturer’s instructions (Illumina).

### MetaRibo-Seq

The lysis supernatant was subjected to ethanol precipitation with 0.1 percent volume 3M sodium acetate and 2.5 volumes of 100 percent ethanol. To precipitate, samples were incubated at -80 °C for 30 minutes, then centrifuged at 21,000 x g for 30 minutes at 4 °C. This was a rough purification specifically implemented to enable suspension of concentrated RNA from reasonable input of fecal sample. The pellet of RNA and RNA-protein complexes was resuspended in MNase buffer. The buffer contained 25 mM Tris pH 8.0, 25 mM NH_4_Cl, 10 mM MgOAc, and 1.55 mM chloramphenicol. To resuspend, we quickly broke the pellet apart with a pipette tip and vortexed for 15 seconds. 1 μL of solution was diluted 20 fold and quantified with Qubit dsDNA HS Assay Kit (Invitrogen). MNase reaction mix was prepared as described^33^, except this was scaled down to an input of 80 μg of RNA and 1 μL of NEB MNase 500 U/μL in a total reaction volume of 200 μL. The MNase reaction was incubated at room temperature for 2 hours. All following steps were performed identically^33^, except the tRNA removal steps were excluded. Briefly, 500 mL of polysome binding buffer was used to wash the Sephacryl S400 MicroSpin columns (GE Healthcare Life Sciences) three times - spinning the column for 3 minutes at 4 °C at 600 RPM. Polysome binding buffer consisted of 100 μL Igepal CA-630, 500 μL magnesium chloride at 1M, 500 μL EGTA at 0.5 M, 500 μL of NaCl at 5M, 500 μL Tris-HCl pH 8.0. at 1M, and 7.9 mL of RNase-free water. The MNase reaction was applied to the column and centrifuged for 5 minutes at 4 °C. The flow through was purified further with miRNAeasy Mini Kit (Qiagen) using manufacture protocols. Elution was performed at 15 μL volume. rRNA was depleted using RiboZero-rRNA Removal Kit for Bacteria (Illumina) using manufacture protocol, except all reaction volumes and amounts were reduced by 50 percent. This was purified with RNAeasy MinElute Cleanup Kit (Qiagen), eluting in 20 uL. The reaction, in 18 μL volume at a total of 100 ng, was subjected to T4 PNK Reaction (NEB M0201S) with addition of 1μL Superase-In (Invitrogen), 2.2 μL 10X T4 PNK Buffer, and 1 μL T4 PNK (10U/μL). This reaction was purified again with RNAeasy MinElute Cleanup (Qiagen). The concentration was determined with Qubit RNA HS Assay Kit (Illumina). With 100 ng as input, libraries were prepared using NEBNext Small RNA Library Prep Set for Illumina (NEB, E7330), using manufacture protocols. DNA was purified using Minelute PCR Purification Kit (Qiagen).

### Small Metatranscriptomics of Fecal Samples

We performed metatranscriptomics as follows: 15 μL of proteinase K (Ambion, 20 mg/mL) was added to 600 μL of lysate. After incubation for 10 minutes at room temperature, samples were centrifuged at 21,000 x g for 3 minutes and the supernatant was collected. An equal volume of Phenol/Chloroform/Isoamyl Alcohol 25:24:1 (pH. 5.2) was applied and vortex for three minutes. The mixture was centrifuged at 21,000 x g for three minutes. The aqueous phase was extracted. This was repeated once more. The final aqueous phase was ethanol precipitated. The RNA was further purified using the RNAeasy Mini plus Kit (Qiagen) using manufacture protocols. Any remaining DNA was degraded via Baseline-ZERO-Dnase (Epicentre) using manufacture protocols. RNA was fragmented for 15 minutes at 70 °C using RNA Fragmentation Reagent (Ambion) using manufacture protocols. At this point, the MetaRibo-Seq and small metatranscriptomics protocol completely converge. The fragmented RNA was purified with miRNAeasy Mini Kit (Qiagen) using manufacture protocols. Elution was performed at 15 μL. rRNA was depleted using RiboZero-rRNA Removal Kit for Bacteria (Illumina) using half reactions of manufacture protocol. This was purified with RNAeasy MinElute Cleanup Kit (Qiagen), eluting in 20 uL. The fragments, in 18 μL volume, were subjected to T4 PNK Reaction (NEB M0201S) with addition of 1μL Superase-In (Invitrogen), 2.2 μL 10X T4 PNK Buffer, and 1 μL T4 PNK (10U/μL). This reaction was purified again with RNAeasy MinElute Cleanup (Qiagen). The concentration was determined with Qubit RNA HS Assay Kit (Invitrogen). With 100 ng as input, libraries were prepared using NEBNext Small RNA Library Prep Set for Illumina (NEB, E7330), using manufacture protocols. DNA was purified using MinElute PCR Purification Kit (Qiagen).

### Differential Centrifugation and FASP for Metaproteomics

To remove human proteins, fecal samples were subjected to differential centrifugation. 100 mg of fecal sample was suspended in 1x PBS in 1.7 mL Eppendorf tubes. The tubes were centrifuged at 600 x g for 1 minute at room temperature. The supernatant was collected in a clean Eppendorf tube and centrifuged at 10,000 x g for 1 minute at room temperature. The supernatant was decanted and the pellet was resuspended in 1 mL of PBS. The process was repeated once more. The final pellet was resuspended in 2% SDS, 100 mM DTT, and 20 mM Tris HCl, pH 8.8 with protease inhibitor. These cells were subjected to bead beating for 3 minutes with a MiniBeadBeater-16, Model 607. 1mM zirconia/silica beads were used. Tubes were centrifuged for 3 minutes and clarified lysate in the supernatant was collected. Lysate was prepared using FASP ^43^ with the same minor modifications previously documented ^44^. Every step involved a centrifugation step for 15 minutes at 14,000 x g. Samples were diluted tenfold in 8 M urea and loaded into Microcon Ultracel YM-30 filtration devices (Millipore). They were washed in 8 M urea, reduced for 30 minutes in 10 mM DTT, and alkylated in 50 mM iodoacetamide for 20 minutes. Samples were washed three times in 8M urea and two times in 50 mM ammonium bicarbonate. Trypsin (Pierce 90057) (1:100 enzyme-to-protein ratio) was added and incubated overnight at 37 °C. Into a new collection tube, samples were centrifuged and further eluded in 50 μL of 70 percent acetonitrile and 1 percent formic acid. The mixture was brought to dryness for one hour using a Savant SPD121P SpeedVac concentration at 30°C, then resuspended in 0.2 percent formic acid^44^.

### Metaproteomics

LC-MS/MS analysis was performed by the Stanford University Mass Spectrometry Facility using the Thermo Orbitrap Fusion Tribrid. A Thermo Scientific Orbitrap Fusion coupled to a nanoAcquity UPLC system (Waters, M Class) was used to collect mass spectra (MS). Samples were loaded on a 25 cm sub 100 micron C18 reverse phase column packed in-house with a 80 minute gradient at a flow rate of 0.45 µL/min. The mobile phase consisted of: A (water containing 0.2% formic acid) and B (acetonitrile containing 0.2% formic acid). A linear gradient elution program was used: 0–45 min, 6–20 % (B); 45-60 min, 35 % (B); 60-70 min, 45 % (B); 70-71 min, 70 % (B); 71-77 min, 95 % (B); 77-80 min, 2 % (B). Ions were generated using electrospray ionization in positive mode at 1.6 kV. MS/MS spectra were obtained using Collision Induced Fragmentation (CID) at a setting of 35 of arbitrary energy. Ions were selected for MS/MS in a data dependent, top 15 format with a 30 second exclusion time. Scan range was set to 400 – 1500 m/z. Typical orbitrap mass accuracy was below 2 ppm; for analysis. A 12 ppm window was allowed for precursor ions and 0.4 Da for the fragment ions for CID fragmentation detected in the ion trap. Prokka-predicted^41^ proteins were used as a reference database for protein detection using the Byonic proteomics search pipeline v 2.10.5^45^. Byonic parameters include: spectrum-level FDR auto, digest cutter C-terminal cutter, peptide termini semi-specific, maximum number of missed cleavages 2, fragmentation type CID low energy, precursor tolerance 12.0 ppm, fragment tolerance 0.4 ppm, protein FDR cutoff 1 percent. These methods were performed by Stanford Mass Spectrometry Facility (SUMS). Using spectral count output, Normalized Spectral Abundance Factor (NSAF) was calculated by in house scripts.

### De Novo Assembly

Quality trimmed metagenomic reads were assembled using metaSPAdes 3.7.0^47^. For all samples, a maximum of 60 million metagenomic reads was used to generate assemblies. Samples sequenced to higher depth were randomly subsetted to 60 million for assembly purposes to both ensure relatively similar numbers of gene predictions and limit computational requirements in assembly and downstream predictions.

### Read Mapping, Gene Prediction and Annotation

Reads were trimmed with trim galore version 0.4.0 using cutadapt 1.8.1^46^ with flags –q 30 and –illumina. Reads were mapped to the annotated assembly using bowtie version 1.1.1^48^. To avoid all possible conservation conflicts in downstream differential analysis, only perfect, unique short read alignments were considered. IGV^49^ was used to visualize coverage. Prokka v1.12^41^ was used to predict genes from the metagenomics assemblies using the –meta option. Annotations were facilitated by many dependencies^38,50,51,52^. For small protein predictions, prodigal^38^ was performed after lowering the size threshold from 90 bases to 60 bases.

### Read density as a function of position

MetaRibo-seq reads were mapped to their metagenomic assemblies. The assembly and aligned reads were analyzed with RiboSeqR^53^. CDSs (coding sequences) were predicted using the findCDS function. Ribosome profiling counts for predicted CDSs were determined with the sliceCounts function. CDSs were filtered to contain at least 10 reads.

### Taxonomic Classification of Technologies

Reads mapping specifically to Prokka-predicted^41^ coding regions were counted. That entire genomic element was input into One Codex^54^ for classification equal to the number of reads mapping to it. This enabled fair comparisons between technologies, as the small metatranscriptomics and MetaRibo-Seq reads can be too small to classify individually with *k*-mer-based approaches. Though metagenomic reads were long enough to be classified directly, they were also subject to the same analysis – entire genes are classified in equal number to the reads overlapping them. Thus, all taxonomy plots represent entire gene classifications and are dependent on the assembly.

### Differential Analysis

The number of reads mapping to a given region were calculated with bedtools multicov^59^. Strandedness was enforced for metatranscriptomics and MetaRibo-Seq. All differential analyses were performed using these counts with all conditions performed in duplicate via DESeq2^60^. A gene was considered differential if it had log2fold change above 1 or below -1, while also reaching an FDR < 0.05. Results were displayed as volcano plots or tables. Heatmaps were created using gplots^61^. Reads per Kilobase Million (RPKM) calculations were performed using in house scripts.

### Statistical Analysis

All Pearson correlations were calculated in R using the Hmisc package^55^. Scatterplots were created with ggplot2^56^. Significance between Pearson correlations was assigned via cocor^57^. Significant differences between RPKM values were assigned using the Kruskal-Wallis test. Significance was assigned as * p value < 0.05, *** p value < 0.001. Zou’s^58^ 95 percent confidence intervals were assigned *** if there is no overlap with 0 in the interval.

### Protein Clustering Analysis

For analyses independent of gene annotation, significantly translationally regulated proteins were clustered using Cd-hit^62^ with 70 percent amino acid identity. Representative sequences were input into Blast2GO^40^ using the nr database. Small protein predictions with translational evidence were also clustered using this same approach. Coding potential was assessed using RNAcode^39^ using the p value assigned to the predicted reading frame.

### Triplet Periodicity Analysis

Using the same default parameters as read density as a function of position, triplet periodicity was called using RiboSeqR^53^. To analyze triplet periodicity of specific genera, assembled contigs were classified using One Codex^54^. Contigs that classified into a specific genus were binned together. Only reads mapping specifically to these bins were considered.

### GO Analysis

Based on differential genes from DESeq2^60^ analyses, UniProt^63^ genes annotated by Prokka^41^ were input into David Functional Annotation^64,65^. All species detected were used as background for these metagenomic analyses.

### Pathway Analysis

Prokka^41^ predicted genes with associated EC (enzyme) numbers were considered. For a given sample, all the reads mapping to any gene with a specific EC number were summed for metatranscriptomics and MetaRibo-Seq. DESeq2^60^ called differential enzymes using MicrobiomeAnalyst^66^. Network mapping is performed to identify pathways corresponding to differential enzymes.

## Supporting information

## Acknowledgements

The authors would like to thank Tessa M Andermann, Ekaterina Tkachenko, and Joyce B. Kang for major contributions to the fecal biobank of bone marrow transplant patients at Stanford hospital. We would like to thank the patients and nurses involved in collection. We thank Christina Wyss-Coray for collecting Alzheimer’s samples used in this study. We appreciate sample collection feedback from Victor Henderson and Tony Wyss-Coray. We thank Anshul Kundaje and Georgi Marinov for helpful computational analysis guidance. We would also like to thank Stanford University Mass Spectrometry for performing and analyzing mass spectrometry on the FASP fecal samples. Sequencing costs were supported via NIH S10 Shared Instrumentation Grant (1S10OD02014101) and Damon Runyon Clinical Investigator Award to ASB, Stanford ADRC grant # P50AG047366. B.J.F is supported by National Science Foundation Graduate Research Fellowship DGE-114747.

---

**Figure S1.**
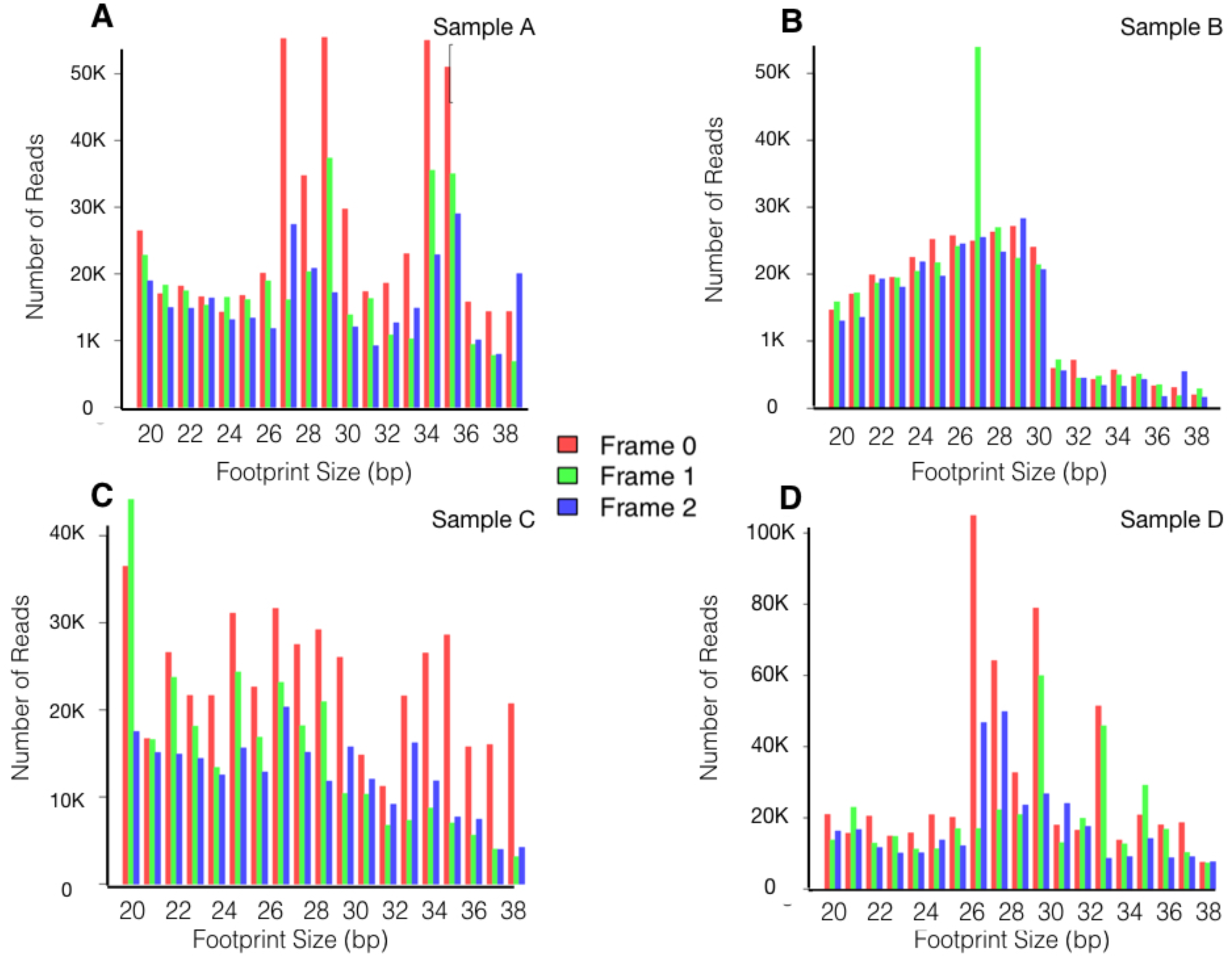
MetaRibo-Seq demonstrates some weak signs of overall codon-resolution. (A-D) Triplet periodicity across footprint lengths for Sample A, B, C, and D, respectively. Colors indicate which frame a read falls within.

**Figure S2.**
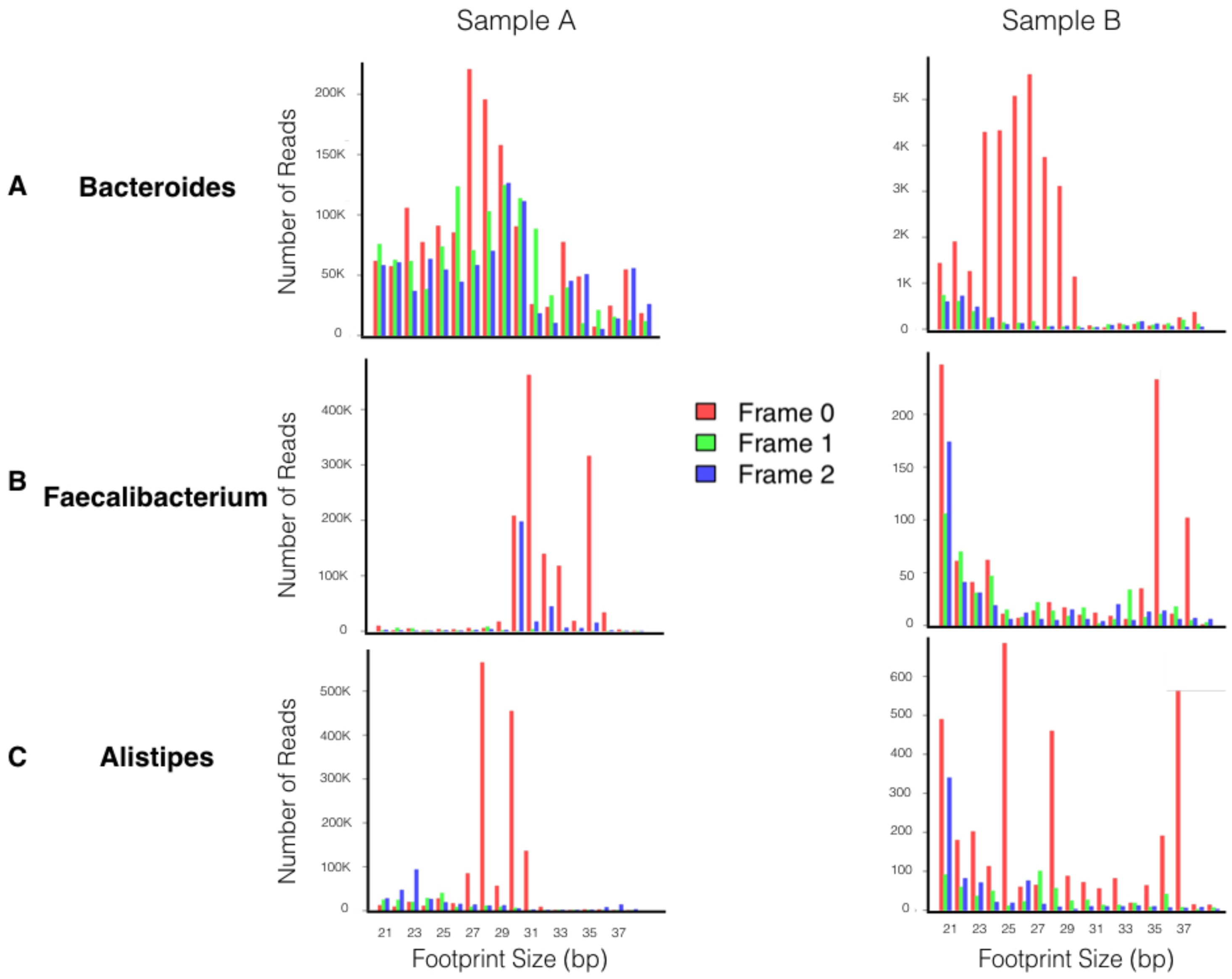
MetaRibo-Seq demonstrates stronger codon resolution in taxa-specific analyses. (A) All contigs assigned to the genera Bacteroides are considered from Sample A and B, respectively. Only these contigs are considered in triplet periodicity analyses. (B and C) The sampe triplet periodicity analysis for Faecalibacterium and Alistipes, respectively.

**Figure S3.**
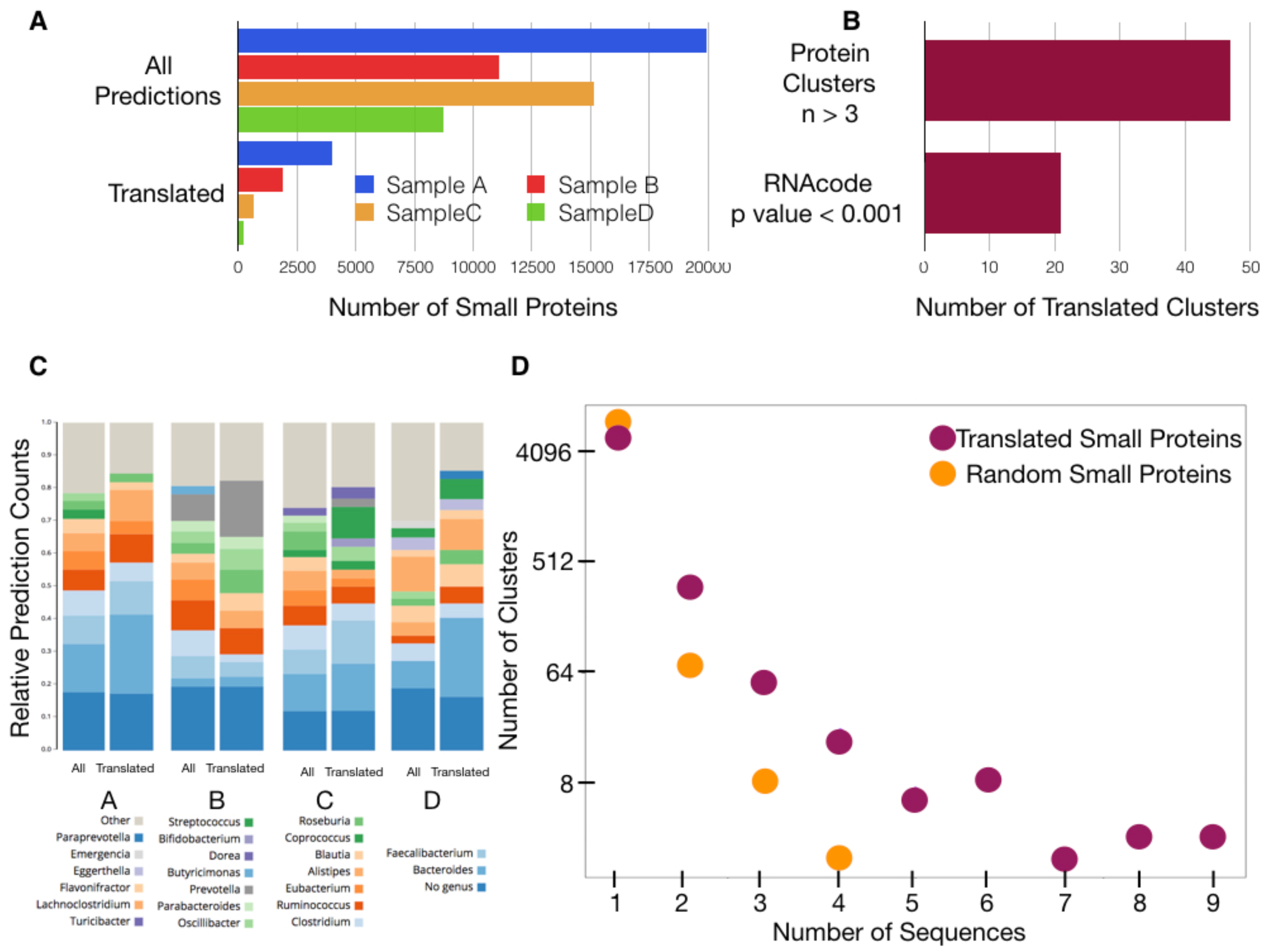
Small protein predictions with translational evidence demonstrate strong conservation. (A) Breakdown of small protein predictions across Samples A, B, C, and D, and those with translational evidence (MetaRibo-Seq signal above 0.5 RPKM). (B) Genus-level classification of small protein predictions across Samples A, B, C, and D. Relative proportions of small proteins including all predictions and only those with translational evidence are provided. (C) The 6,774 small proteins with translational evidence are clustered at 70 percent amino acid identity. 47 clusters with at least 4 members are identified. Among these, 21 clusters possess evolutionary signatures indicative of coding regions (p values < 0.001) via RNAcode^39^. (D) In dark red, clustering with 70 percent protein identity of small proteins with translational evidence – 6,774 proteins across all samples. In orange, clustering with 70 percent amino acid identity of 6,744 small proteins randomly chosen from prodigal predictions; an equal number of proteins as those with translational evidence from each sample were randomly chosen.

**Figure S4.**
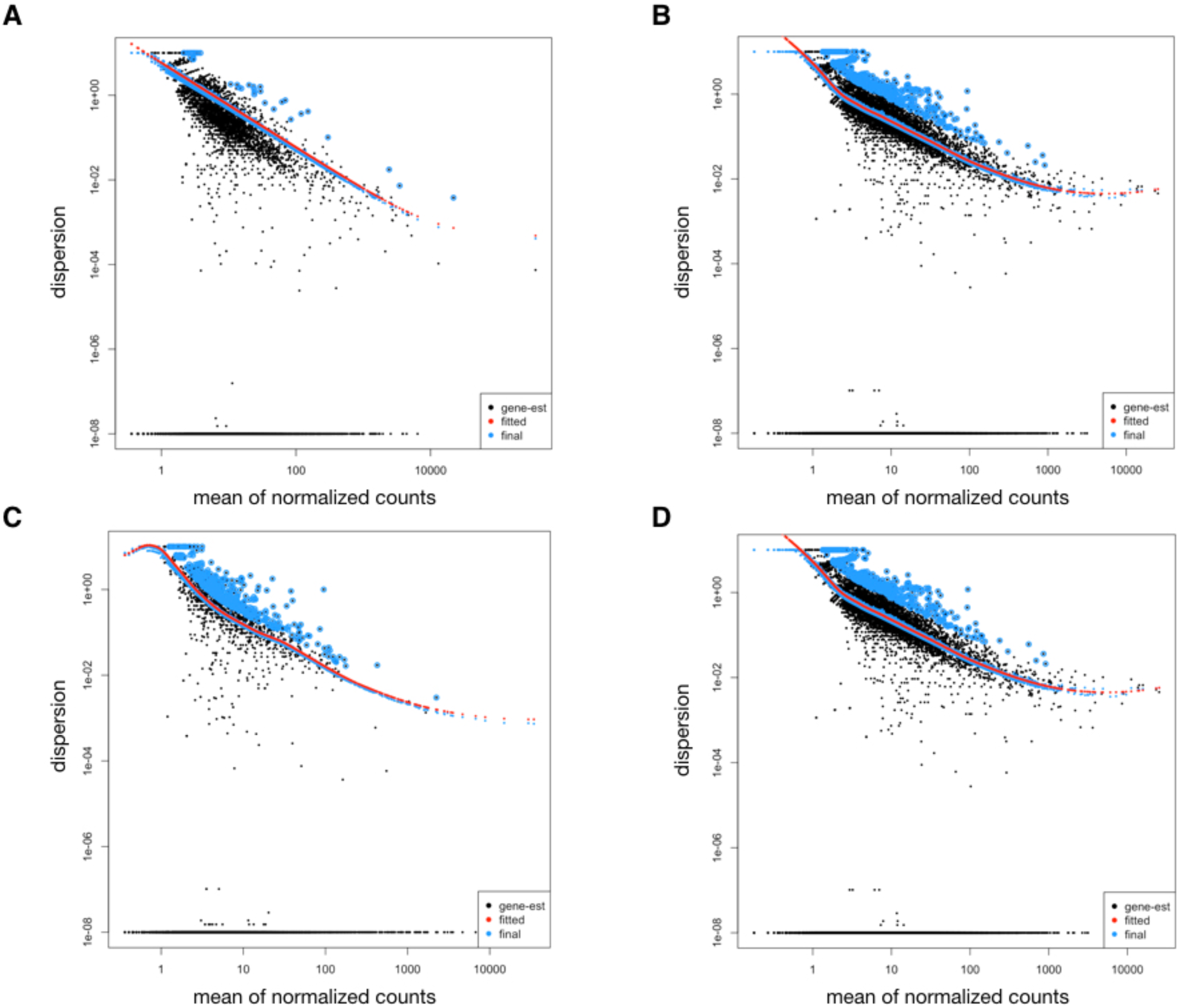
DESeq2 models adequately fit dispersion in comparing metatranscriptomics to MetaRibo-Seq. (A-D) Dispersion plots of DESeq2 models fit to Samples A, B, C, and D, respectively.

**Figure S5.**
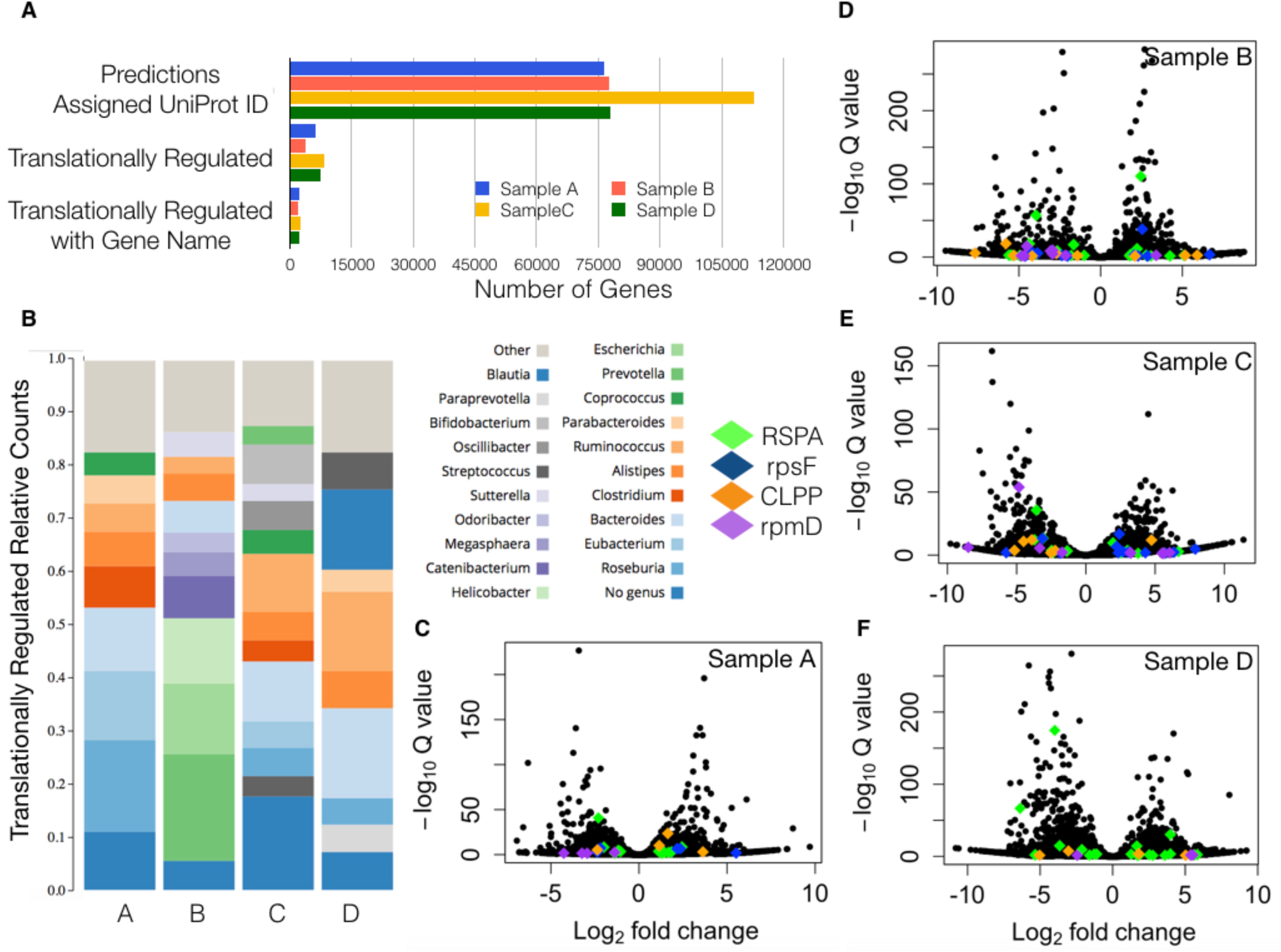
Numerous genes are consistently translationally regulated in the fecal microbiota. (A) For Samples A, B, C and D, we show genera level classifications of translationally regulated genes assigned UniProt^31^ protein IDs. (B-E) Volcano plots comparing metatranscriptomics and MetaRibo-Seq in Sample A, B, C, and D, respectively. Significant genes are colored in red. The four most consistently translationally regulated genes are also denoted.

**Figure S6.**
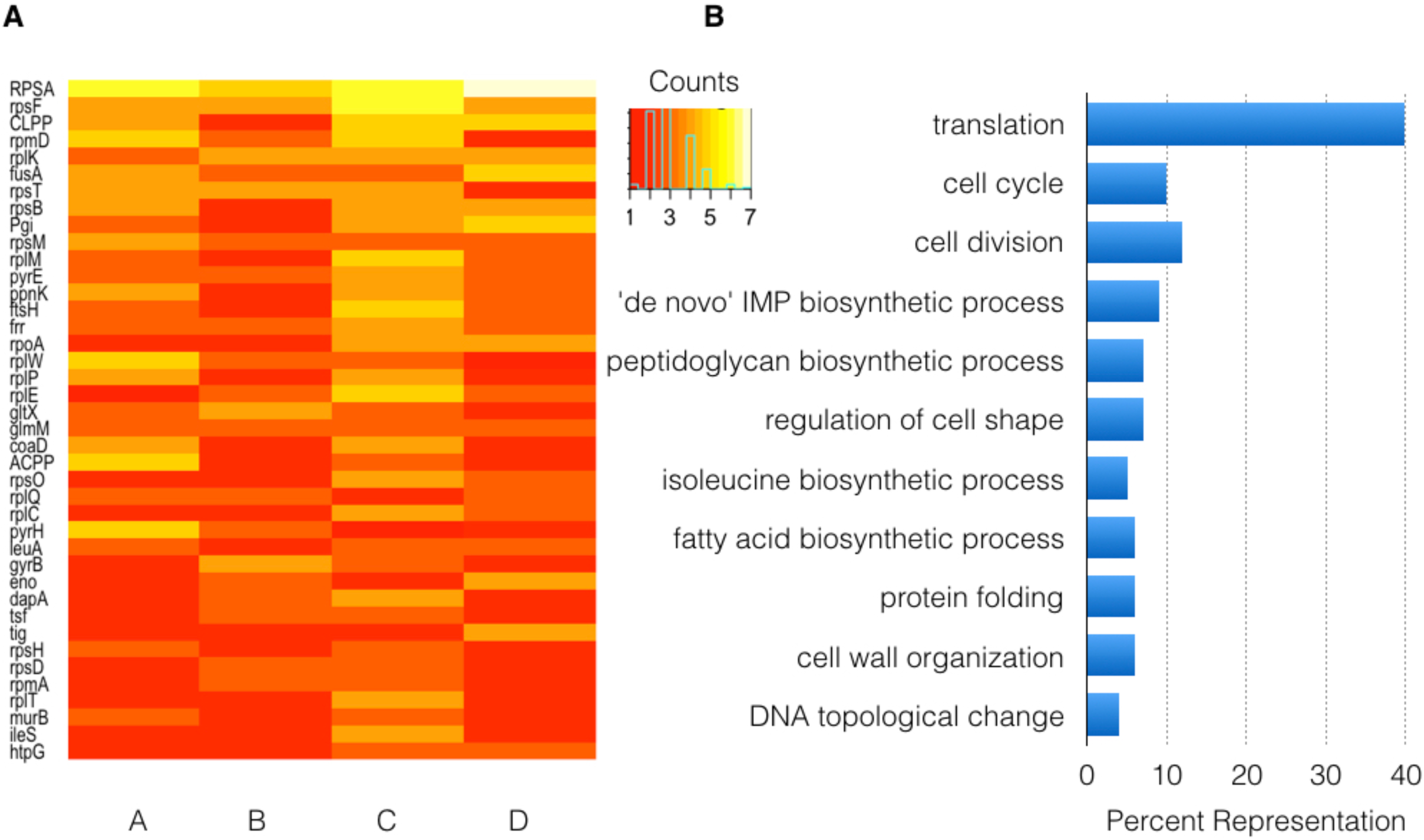
In depth analysis of consistently translationally regulated genes with assigned names in the fecal microbiota across Samples A, B, C, and D. (A) Top 40 most common translationally regulated genes across samples. Heatmap intensity represents the number of times the gene appeared differential in a sample. (B) GO analysis including genes that are differentially translated at least 5 times across samples.

**Table S1**

Mapping statistics to *de novo* references. This table shows the number of reads, percentage of reads mapping to the assembly, and percentage of reads overlapping regions annotated as coding.

**Table S2**

GO analysis of significant translationally regulated genes for Samples A, B, C, and D. Only genes assigned UniProt^63^ protein IDs are considered. Significant IDs are called with DESeq2^60^. These are input into David^64^.

**Table S3**

Pairwise comparison of metatranscriptomics to MetaRibo-Seq across Sample A, B, C, and D. Only predicted genes with EC numbers are considered. Significant differences in EC numbers are called with DESeq2^60^. Negative log2FC means lower in the technology first listed in the tab under consideration. EC numbers and their associated adjusted p value are input into MicrobiomeAnalyst^66^ to determine overrepresented pathways based on differential EC numbers.

**Table S4**

We display all correlations between replicates and technologies for Samples A, B, C, and D. For each sample, we provide correlations between metatranscriptomics replicates, MetaRibo-Seq replicates, metagenomics versus metatrancriptomics, metagenomics versus MetaRibo-Seq, metatranscriptomics versus MetaRibo-Seq, and metagenomics versus translation efficiency (TE – ratio of MetaRibo-Seq and metatranscriptomics).

**File S1**

We provide sequences for every protein in Samples A, B, C, and D that are translationally regulated in the gut microbiota. Translationally regulated proteins (.faa) and 70 percent identity clustering (.cltsr) are provided for Samples A, B, C, and D. The representative sequences for clusters (.faa) with more than 5 sequences for samples are also provided. The sequence name itself denotes which sample the sequence is found in. Any sequence that begins with a specific identifier can be linked to a sample: HDALDHFB = Sample A, HENMDNCI =Sample B, PJJNKMKO = Sample C, GPBGFMPE = Sample D. Blast2GO^40^ results for the representative sequences of consistent clusters are provided.

**File S2**

We contribute sequences of small proteins we identified. Small protein predictions using prodigal^38^ with lower size threshold (60 bp) for Samples A, B, C, and D individually (.faa) are provided. Small protein predictions with MetaRibo-Seq RPKM above 0.5 for the four samples individually (.faa) are given. Combined clustering of these small proteins with translational evidence are provided (.clstr). Those translated clusters with more than 3 sequences and with RNAcode p values < 0.001 are provided in the “smallproteinsequences” folder named by the cluster they belong to.

